# A Toxic RNA Catalyzes the Cellular Synthesis of Its Own Inhibitor, Shunting It to Endogenous Decay Pathways

**DOI:** 10.1101/741926

**Authors:** Raphael I. Benhamou, Alicia J. Angelbello, Eric T. Wang, Matthew D. Disney

## Abstract

Myotonic dystrophy type 2 (DM2) is a genetically defined muscular dystrophy caused by a toxic expanded repeat of r(CCUG) [heretofore (CCUG)^exp^], harbored in intron 1 of CHC-Type Zinc Finger Nucleic Acid Binding Protein (*CNBP*) pre-mRNA. This r(CCUG)^exp^ causes DM2 via a gain-of-function mechanism that results in three hallmarks of its pathology: (i) binding to RNA-binding proteins (RBPs) that aggregate into nuclear foci; (ii) sequestration of muscleblind-like-1 (MBNL1) protein, a regulator of alternative pre-mRNA splicing, leading to splicing defects; and (iii) retention of intron 1 in the *CNBP* mRNA. Here, we find that *CNBP* intron retention is caused by the r(CCUG)^exp^-MBNL1 complex and can be rescued by small molecules. We studied two types of small molecules with different modes of action, ones that simply bind and ones that can be synthesized by a r(CCUG)^exp^-templated reaction in cells, that is the RNA synthesizes its own drug. Indeed, our studies completed in DM2 patient-derived fibroblasts show that the compounds disrupt the r(CCUG)^exp^-MBNL1 complex, reduce intron retention, subjecting the liberated intronic r(CCUG)^exp^ to native decay pathways, and rescue other DM2-associated cellular defects. Collectively, this study shows that small molecules can affect RNA biology by shunting toxic transcripts towards native decay pathways.

**HIGHLIGHTS:**

- Intron retention in RNA repeat expansions can be due to repeats binding to proteins
- Small molecules that bind RNA repeats and inhibit protein binding can trigger decay
- A toxic RNA repeat can catalyze the synthesis of its own inhibitor on-site
- On-site drug synthesis most potently affects disease biology

**eTOC BLURB:** The most common way to target RNA is to use antisense oligonucleotides to target unstructured RNAs for destruction. Here, we show for the first time that small molecules targeting structured, disease-causing RNAs can shunt them towards native decay pathways by affecting their processing.

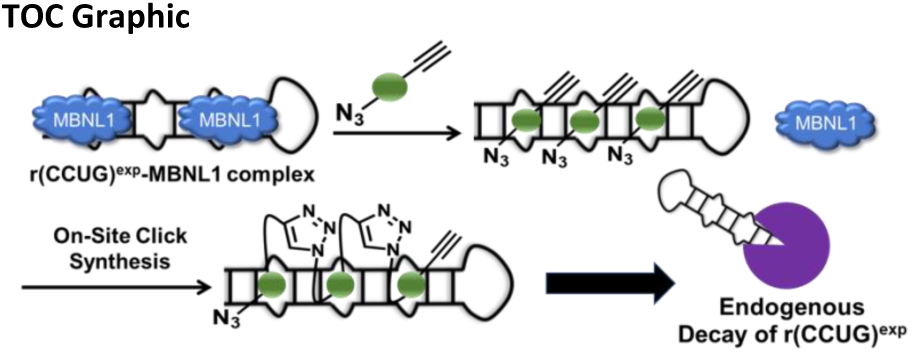

## INTRODUCTION

A variety of human diseases are mediated by RNA structures [1-3], ideal targets for small molecules that could elucidate RNA biology or be exploited for therapeutic development [4-6]. Estimates suggest that the number of potential human RNA drug targets are an order of magnitude greater than the number of protein targets [6, 7]. Compounds that bind RNA have been identified by using screening [8-10], structure-based design [11, 12], or sequence-based design [13]. However, only a very limited number of these small molecules are bioactive and far fewer are known to target human RNAs and affect their function [4]. As these tools are more fully developed and more broadly deployed, they will likely deliver many chemical probes to study human RNA biology.

RNA repeating transcripts are attractive targets for chemical probes, causing >30 genetically defined diseases with no known treatments [14]. Although the RNA is the central toxin in each disease, their pathomechanisms and impact on biological processes are diverse. For example, repeats can cause transcriptional silencing by binding to promoters from which they are encoded, as has been observed in fragile X syndrome (FXS) [15]. Toxic RNAs can be translated independently of a canonical start codon, in a process named repeat associated non-ATG (RAN) translation, as observed in C9orf72 amyotrophic lateral sclerosis and frontotemporal dementia (c9ALS/FTD) [16]. RNA repeats also cause toxicity by a gain-of-function mechanism in which they bind to and sequester proteins involved in RNA biogenesis, forming nuclear foci and causing defects in RNA processing, as observed in myotonic dystrophy types 1 and 2 (DM1 and DM2, respectively) [17, 18]. More recently, various RNA repeats have been shown to affect the processing of the pre-mRNAs in which they reside, causing intron retention and providing potential biomarkers [19].

DM2 is caused by an expanded r(CCUG) repeat, or r(CCUG)^exp^, located in intron 1 of the CHC-Type Zinc Finger Nucleic Acid Binding Protein (*CNBP*) pre-mRNA. Like other RNA repeat expansions, the DM2 RNA is highly structured and forms a periodic array of 5’C<u>CU</u>G/3’G<u>UC</u>C 2×2 internal loops (Figure 1A). We previously designed two types of small molecules that recognize this repeating structure and improve DM2-associated defects [20-22]. In particular, they rescue the aberrant splicing of bridging integrator 1 (*BIN1*) exon 11 and the formation of nuclear foci in a mouse myoblast cell line transfected with a plasmid encoding r(CCUG)_300_ [21, 22]. The first compound, a dimer comprised of two RNA-binding modules, recognizes and binds two adjacent loops simultaneously, which affords avidity and selectivity [21]. The second class of compounds are synthesized *in cellulis* and only in DM2-affected cells [22]; that is, r(CCUG)^exp^ synthesizes its own inhibitor. In cell, on-site drug synthesis is enabled by appending the RNA-binding modules that target the 2×2 internal loops in r(CCUG)^exp^ with azide and alkyne units that are unreactive in biological systems unless brought into high effective concentration or proximity by binding adjacent sites in r(CCUG)^exp^ [22].

**Figure 1:**
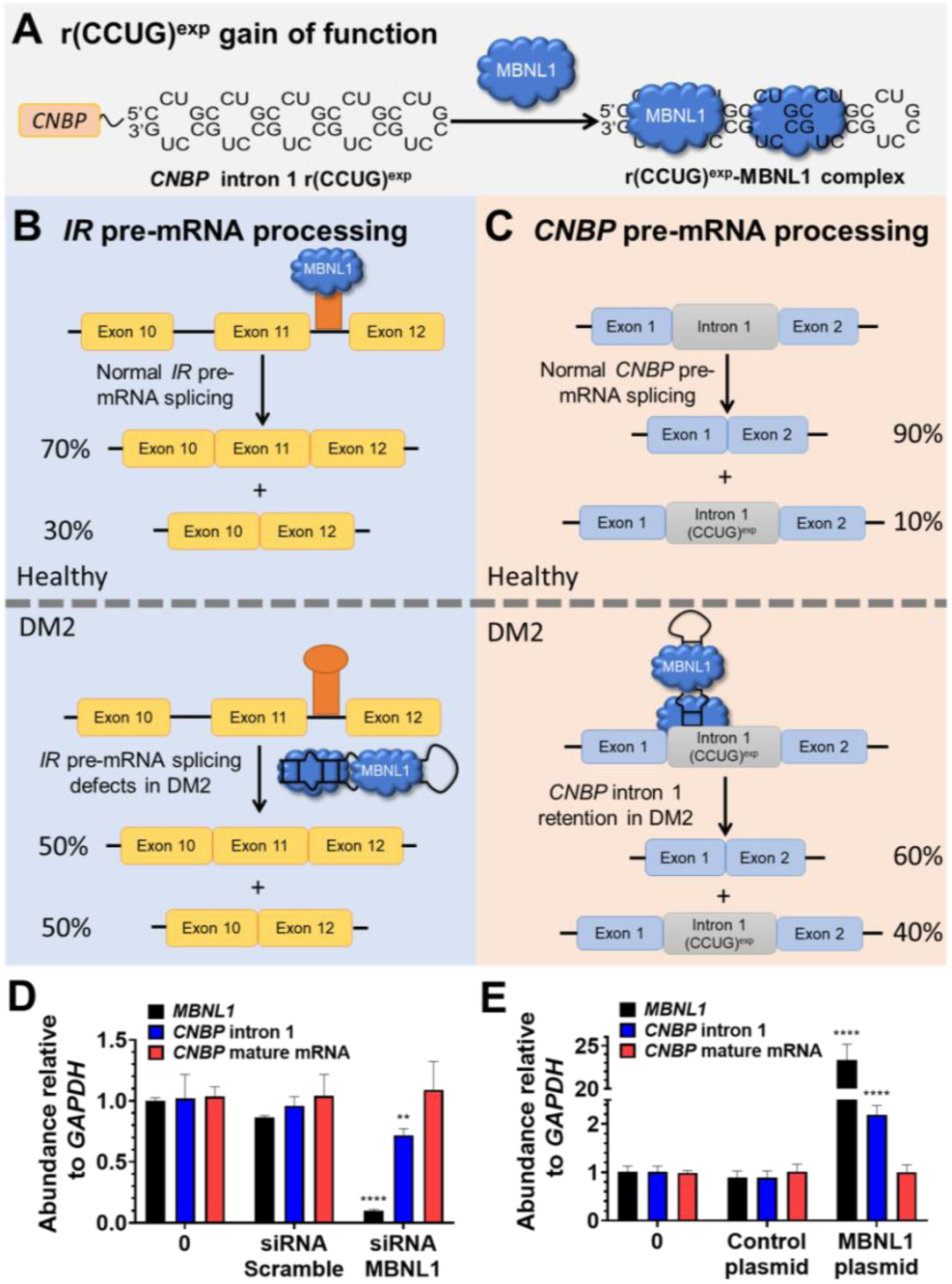
The RNA that causes DM2, r(CCUG)^exp^, has multiple modes of toxicity. (*A*) r(CCUG)^exp^ in intron 1 of *CNBP* folds into a hairpin displaying a periodic array of internal loops that sequester MBNL1. Sequestration of MBNL1 by r(CCUG)^exp^ causes (*B*) pre-mRNA splicing defects, for example increasing *IR* exon 11 exclusion and (*C*) aberrant retention of *CNBP* intron 1. (*D*) MBNL1 knock-out using siRNA and its effect on relative abundance of *MBNL1, CNBP* intron 1, and *CNBP* mature mRNA as measured by RT-qPCR. (*E*) MBNL1 knock-in using a transfected MBNL1 plasmid and its effect on relative abundance of *MBNL1, CNBP* intron 1, and *CNBP* mature mRNA as measured by RT-qPCR. Error bars represent SD. \*\**P* < 0.01, \*\*\**P* < 0.001, \*\*\*\**P* < 0.0001 as determined by a one-way ANOVA (n = 3).

Herein, we have evaluated these compounds in DM2 patient-derived fibroblasts. Importantly, these studies revealed that intron retention in DM2-affected cells is caused by MBNL1 and that this defect can be rescued by small molecules, a pathway only defined by studying the RNA in its native context. That is, small molecules that prevent or disrupt MBNL1 binding not only alleviate pre-mRNA splicing defects and reduce nuclear foci, but they also rescue *CNBP* intron retention, shunting the r(CCUG)^exp^-containing intron to RNA quality control pathways and eliminating the toxin.

## RESULTS

### Intron retention in DM2 patient-derived fibroblasts

As aforementioned, r(CCUG)^exp^ has at least three pathomechanisms: (i) formation of nuclear foci when it binds various RBPs; (ii) pre-mRNA splicing defects in MBNL1-regulated genes [20, 21, 23, 24]; and, (iii) retention of intron 1 in *CNBP* mRNA, in which it is harbored [19] (Figure 1). The first two mechanisms were identified shortly after the discovery of the repeat expansion, facilitated by in-depth studies completed for the repeat that causes DM1. In contrast, little is known about the mechanism of intron retention. With chemical probes that bind r(CCUG)^exp^ in hand, we sought to study if *CNBP* intron 1 retention was also stimulated by MBNL1 binding to r(CCUG)^exp^ (Figure 1C).

To study the role of MBNL1 in these events, a series of gain-and loss-of-function experiments were completed in DM2 patient-derived fibroblasts. When a siRNA directed at *MBNL1* was delivered to DM2 fibroblasts, a significant decrease in intron 1 retention was observed (Figures 1D and S1). In contrast, no effect was observed when cells were treated with a control, scrambled siRNA (Figure S1).

To further support the role of MBNL1 in intron retention, a plasmid encoding *MBNL1* was transfected into DM2 fibroblasts. An increase in the amount of intron retained product was observed upon overexpression of MBNL1 but not with a control plasmid (Figure 1E and Figure S1). Collectively, both studies support a role for MBNL1 binding to r(CCUG)^exp^ in the formation of intron retained products.

### A designed dimeric compound (**1**) that targets the structure of r(CCUG)^exp^ reduces nuclear foci and rescues alternative splicing

Previously, a series of monomeric and dimeric ligands were designed that target the structure of r(CCUG)^exp^ (Figure 2A) [21, 25]. These compounds are built on a kanamycin A scaffold in which the 6’ position is acylated. The consequence of this modification is that the compounds bind specifically to the internal loops in r(CCUG)^exp^ and have no measurable affinity to the A-site in the bacterial ribosome [20, 21, 23], a target of aminoglycosides [24]. Based on these previous studies, a compound with two kanamycin modules, **1**, was designed that binds two 5’C<u>CU</u>G/3’G<u>UC</u>C internal loops simultaneously (Figure 2B). The compound binds to r(CCUG)^exp^ with low nanomolar affinity and selectively discriminates against a number of other RNA targets [20, 26]. Further, **1** rescued DM2-associated splicing defects and reduced the number of nuclear foci in a transfected model system, as DM2 patient-derived cells were not available [21]. Studies on intron retention were not possible in this cellular model as only the repeats themselves were expressed, outside the context of the *CNBP* transcript.

**Figure 2:**
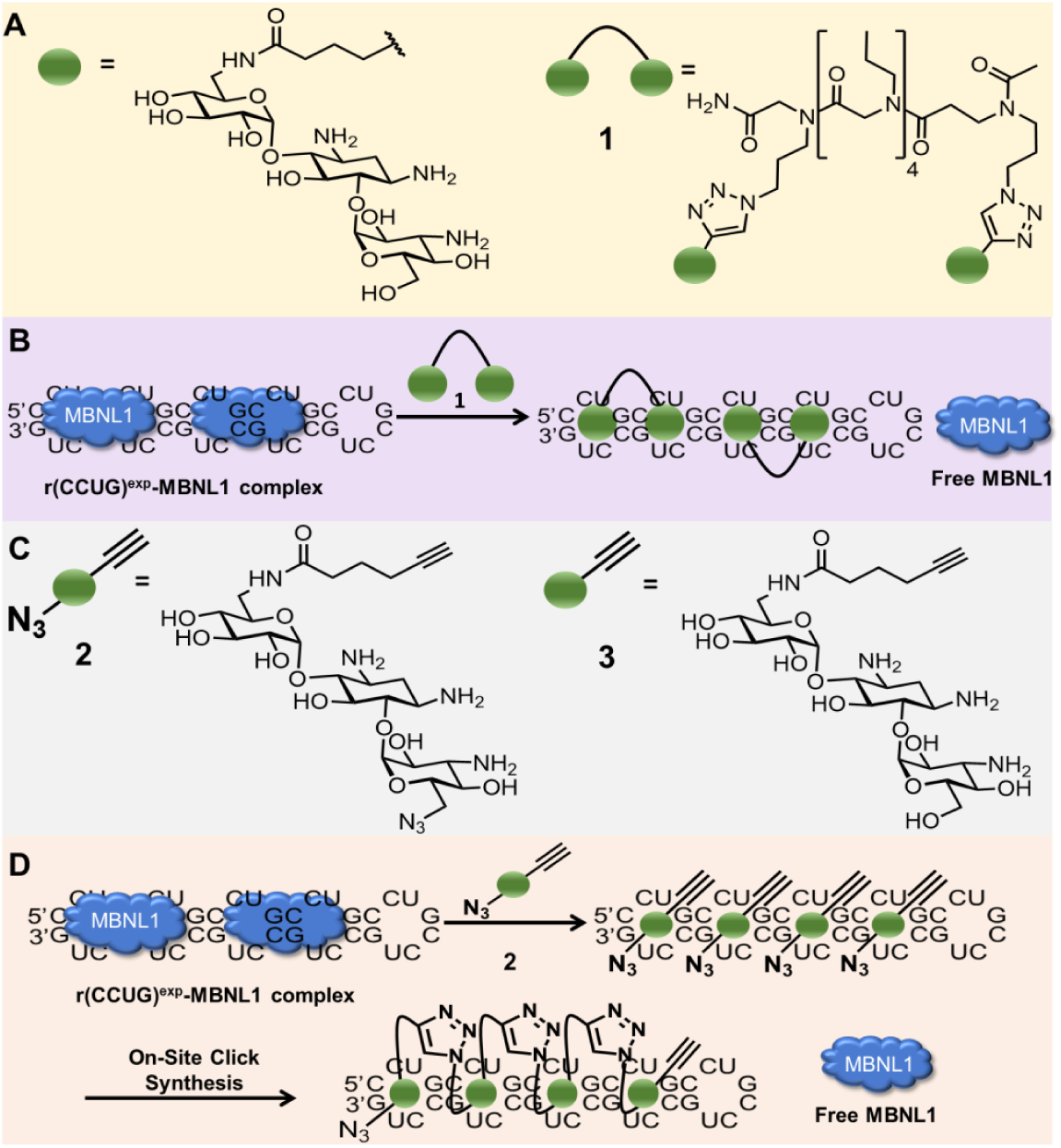
Design and biological impact of small molecules targeting r(CCUG)^exp^. (*A*) Structures of a kanamycin derivative (green spheres) that avidly binds the internal loops found in r(CCUG)^exp^ and a dimer composed of two kanamycin binding modules connected via a propylamine peptoid (**1**). (*B*) Schematic of **1**’s mode of action, binding to r(CCUG)^exp^ and releasing MBNL1. (*C*) Structures of compounds employed in an on-site drug synthesis approach. Compound **2**, a modified kanamycin module, contains alkyne and azide moieties that react upon binding r(CCUG)^exp^, the catalyst. Compound **3** is a kanamycin module containing only the alkyne component and thus cannot oligomerize. (*D*) Schematic of on-site click synthesis of **2**, catalyzed by binding to r(CCUG)^exp^, and release MBNL1 to relieve DM2-associated defects.

We therefore extended those studies to DM2 patient-derived cells, investigating **1**’s ability to rescue formation of nuclear foci, pre-mRNA splicing defects, and importantly intron retention. Indeed, treatment with **1** reduced formation of nuclear foci, with a statistically significant effect observed at 10 μM (P <0.01; Figure 3A&B). Notably, both the presence of r(CCUG)^exp^ and MBNL1 in foci were assessed in our analysis, indicating liberation of MBNL1 from r(CCUG)^exp^.

**Figure 3:**
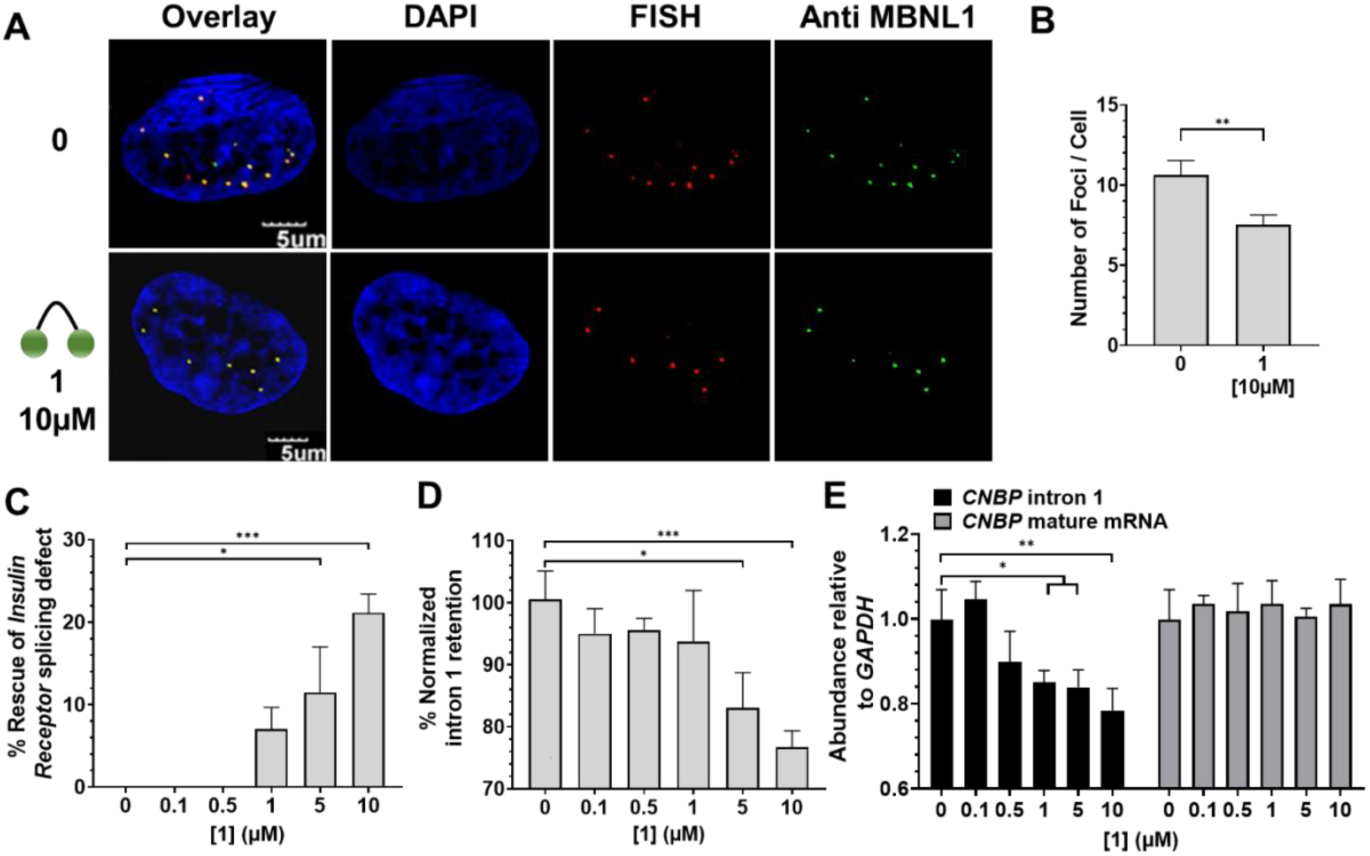
Biological activity of **1** in DM2 patient-derived fibroblasts. (*A*) Representative microscopic images for the reduction of nuclear foci by **1**, as completed by RNA-FISH and immunohistochemistry using an anti-MBNL1 antibody. (*B*) Quantification of r(CCUG)^exp^-MBNL1 foci/nucleus (n = 3 biological replicates, 40 nuclei counted per replicate). *(C)* Effect of **1** on an MBNL1-regulated alternative splicing event, *IR* exon 11, as determined by RT-PCR; *(D)* Effect of **1** on aberrant *CNBP* pre-mRNA splicing caused by r(CCUG)^exp^, as determined by RT-PCR; that is, retention of intron 1; *(E)* Effect of **1** on total intron 1 levels and *CNBP* mature mRNA, as determined by RT-qPCR. Error bars represent SD. \**P* < 0.5, \*\**P* < 0.01, \*\*\**P* < 0.001, \*\*\*\**P* < 0.0001 as determined by a one-way ANOVA (n = 3).

The alternative splicing of insulin receptor (*IR*) exon 11 is controlled by MBNL1; exon 11 is excluded too frequently in DM2 (Figure 1B) [27]. Compound **1** improved *IR* exon 11 alternative splicing defects dose-dependently while not affecting splicing in WT fibroblasts (Figures 3C, S2 & S3). Additionally, no effect was observed on the alterative splicing of mitogen-activated protein kinase kinase kinase kinase 4 (*MAP4K4*) exon 22a, which is not MBNL1-dependent (Figure S4).

### Designer compounds that target the structure of r(CCUG)^exp^ decrease intron retention

Two lines of experimental evidence suggest that **1** liberates a fraction of MBNL1 such that it can resume its normal function: (i) reduced nuclear foci containing both r(CCUG)^exp^ and MBNL1 and (ii) improvement of a MBNL1-dependent splicing defect. We therefore hypothesized that **1** might also rescue MBNL1-dependent intron retention. Indeed, **1** reduced the amount of intron 1 retained in *CNBP* mRNA in DM2 fibroblasts as measured by RT-PCR, indicating that the small molecule rescues aberrant *CNBP* pre-mRNA splicing (Figure 1C, 3D, and S5).

Intronic regions are subjected to decay when liberated from their pre-mRNAs [28]. Thus, we tested the possibility that **1** could stimulate endogenous decay of the freed intron containing r(CCUG)^exp^[28]. By using RT-qPCR primers for intron 1, we found that ∼20% of the intron is eliminated from cells (Figure 3E). Importantly, **1** has no effect on *CNBP* mature mRNA levels (Figure 3E & S5) and did not affect the abundance of intron 1 in healthy cells (Figure S6). Thus, binding of the ligand affects the processing of the pre-mRNA in a manner that then eliminates the toxic repeat. Collectively, these data further support the hypothesis that MBNL1 sequestration stimulates intron 1 retention of *CNBP* while simultaneously demonstrating a new activity for small molecules that bind RNA targets, subjecting the RNA to quality control pathways to facilitate its degradation.

### The toxic r(CCUG)^exp^ synthesizes its own inhibitor in patient-derived cells

Although binding compound **1** rescued three hallmarks of DM2, its activity is not as robust as desired. We previously reported an approach to synthesize oligomeric compounds with enhanced potency (∼100-fold more potent than the simple binding compound) and selectivity in a cell line transfected with r(CCUG)_300_ [22]. The approach, inspired by Sharpless to use click chemistry to assemble compounds on acetylcholinesterase *in vitro* [29], was also used by the Dervan group to assemble polyamides on a DNA minor groove *in vitro* [30]. We demonstrated that RNA repeats can template the synthesis of their own inhibitors using a click reaction *in cellulis*, including in DM1 patient-derived cells [22, 31]. Notably, the clickable compounds developed for DM1 and DM2 are the most potent RNA-targeting small molecules identified to date with pM and nM EC_50_’s for rescuing disease-associated defects in cells, respectively, and are 50,000-fold more potent than the corresponding repeat-targeting oligonucleotides [31, 32].

Briefly, for r(CCUG)^exp^-catalyzed on-site, in cell drug synthesis, a kanamycin moiety was appended with biorthogonal azide and alkyne moieties, affording **2**. When **2** binds adjacent 2×2 nucleotide internal loops in r(CCUG)^exp^, the azide displayed by one **2** molecule is brought into close proximity with the alkyne of another **2** molecule, reacting to form stable triazole units and hence multimeric compounds. This reaction is catalyzed by the RNA and does not require copper(I). We therefore studied if on-site drug synthesis using **2** is also operational in DM2 fibroblasts and occurs selectively; that is, on-site synthesis should not occur in fibroblasts lacking r(CCUG)^exp^ (i.e., WT). For these studies, DM2 and WT fibroblasts were co-treated with **2** and **3**, a kanamycin derivative that only contains an alkyne moiety. Compound **3** limits the extent of polymerization, thereby allowing mass spectrometry to be used to assess reactivity in cells. Indeed, mass spectral analysis of lysates from treated DM2 fibroblasts indicated formation of both trimer and tetramer products, in agreement with our previously reported studies in transfected cells (Figure S7) [22]. Importantly, oligomerization was not observed in healthy fibroblast treated with **2** and **3**, since no r(CCUG)^exp^ is present to catalyze the click reaction (Figure S7).

### In cell synthesized compounds reduce foci and rescue splicing defects more potently than **1**

To compare the potency of **1** to compounds synthesized on-site derived from **2**, we evaluated the ability of **2** to reduce the number of r(CCUG)^exp^-MBNL1 foci in cells. On average, DM2 fibroblasts contain 10.6 ± 0.9 foci per cell. Upon treatment with 1 μM of **2**, the number of foci was reduced by ∼45% (5.7± 0.9) (Figure 4A&B), a significant improvement over **1**, which reduced the number of foci by only ∼20% at 10 μM concentration (Figure 3A&B). As a control, DM2 fibroblasts were treated with **3**, which cannot oligomerize. No significant decrease in foci was observed (Figure S8).

**Figure 4:**
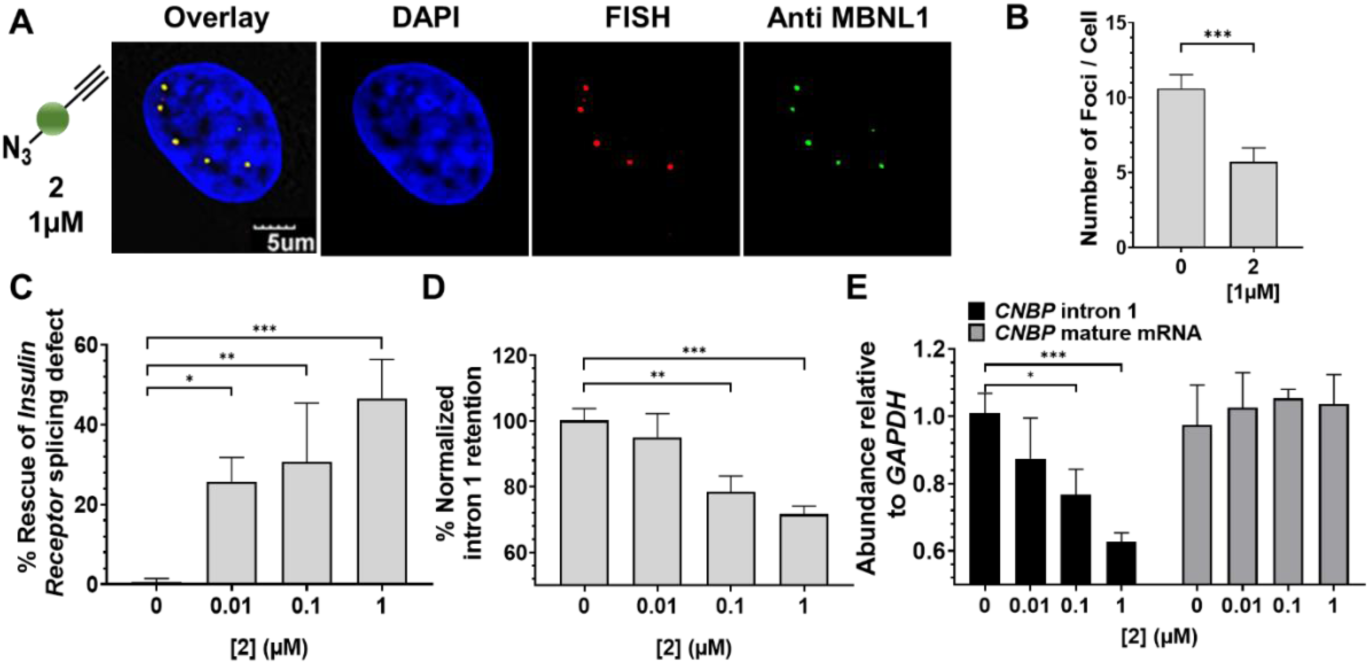
Biological activity of **2** in DM2 patient-derived fibroblasts. (*A*) Representative microscopic images for the reduction of nuclear foci by **2**, as completed by RNA-FISH and immunohistochemistry using an anti-MBNL1 antibody. (*B*) Quantification of r(CCUG)^exp^-MBNL1 foci/nucleus (n = 3 biological replicates, 40 nuclei counted per replicate). *(C)* Effect of **2** on an MBNL1-regulated alternative splicing event, *IR* exon 11, as determined by RT-PCR; *(D)* Effect of **2** on aberrant *CNBP* pre-mRNA splicing caused by r(CCUG)^exp^, as determined by RT-PCR; that is, retention of intron 1; *(E)* Effect of **2** on total intron 1 levels and *CNBP* mature mRNA, as determined by RT-qPCR. Error bars represent SD. \**P* < 0.5, \*\**P* < 0.01, \*\*\**P* < 0.001 as determined by a one-way ANOVA (n = 3).

Given these encouraging results, we tested the ability of **2** to correct the *IR* alternative pre-mRNA splicing defect (Figures 1B & 4C). Notably, ∼45% rescue of *IR* exon 11 splicing was observed at 1 µM of **2** (Figure 4C & S9), compared to only 20% rescue with the dimer **1** at 10 µM (Figure 3C). Control compound **3** showed no significant rescue of the *IR* splicing defects in DM2 fibroblasts, as expected (Figure S10). Additionally, **2** does not affect *IR* alternative splicing in healthy fibroblasts (Figure S11) nor splicing of *MAP4K4* exon 22a in DM2 fibroblasts (Figure S12).

### In cell synthesized compounds derived from **2** decrease intron retention more potently than **1**

To further confirm that compounds derived from the oligomerization of **2** liberate MBNL1 from r(CCUG)^exp^ to improve DM2 pathology, we studied its ability to rescue the aberrant splicing of *CNBP* pre-mRNA and to activate the cellular decay of the liberated intron containing r(CCUG)^exp^, as observed for **1**. Clickable compound **2** reduced the amount of intron 1 retained in *CNBP* mRNA dose dependently, with an ∼30% reduction observed upon treatment with 1 μM (Figure 4D), an ∼10-fold improvement over **1** (Figure 3D & S9). Likewise, using RT-qPCR primers for intron 1 itself, **2** reduced overall intron 1 levels dose-dependently with an ∼40% reduction at 1 μM (Figure 4E). For comparison, **1** only reduced intron 1 levels by ∼20% when DM2 fibroblasts were treated with 10 μM compound. Importantly, this reduction can be traced directly to rescue of aberrant *CNBP* splicing as levels of mature *CNBP* mRNA were not affected (Figure 4E). Further, **2** did not affect the abundance of intron 1 in healthy cells (Figure S13). As expected, **3** is inactive; it does not affect intron 1 levels in DM2 fibroblasts (Figure S14) nor does it affect *CNBP* intron 1 levels or *IR* splicing in healthy fibroblasts. Thus, on-site drug synthesis converts an inactive compound into a potent modulator of function.

## DISCUSSION

Two significant challenge have precluded exploiting RNA as a drug target: (i) the lack of approaches and rules that govern their molecular recognition with non-oligonucleotide-based modalities; and (ii) a companion set of data that defines which RNA-small molecule binding events elicit a biological response [33]. Various tools have emerged to study molecular recognition events and to validate RNA targets *in vitro* and *in cellulis*, which have begun to provide these much needed data [4]. Such tools include small molecules with novel activities *in situ* and *in vivo* that directly and selectively cleave the RNA target [34] or recruit cellular nucleases to effect their degradation [35].

Of particular interest, this study has shown that a toxic RNA can template the synthesis of its own inhibitor in patient-derived cells to rescue multiple disease mechanisms. The observation that **2** can be transformed into a bioactive oligomeric species upon binding to adjacent sites in r(CCUG)^exp^ suggests that disease-affected cells can be provoked to manufacture a cellular active inhibitor from an inactive monomer (**3** in this case) at the needed site of action. Healthy cells that have the same genetic makeup but do not express the disease-causing RNA target were therefore unexposed to the active compounds, as oligomerization did not occur. Many microsatellite disorders, such as c9ALS/FTD, are caused by RNAs that are expressed in the brain. An on-site synthesis approach could therefore be advantageous, providing low molecular weight, tissue and cell penetrant compounds that can engage the disease-causing targets and be assembled into potent modulators of RNA function *in situ*.

A second aspect of special interest is the observation that RNA repeat expansions harbored in introns may be especially sensitive to small molecule intervention, the activity of which is then interfaced with endogenous RNA quality control decay mechanisms. A variety of other RNA repeating transcripts are present in introns and form intron retained products. A previous study showed that GC-rich repeats harbored in introns [such as r(CCUG)^exp^] cause the greatest extent of retention (as compared to AU-rich repeats), indicating a potential role for protein loading in intron retention observed in each disease [19]. These GC-rich repeat expansions, which cause c9ALS/FTD [r(G_4_C_2_)^exp^] repeat expansions, Fuchs endothelial corneal dystrophy [FECD; r(CUG)^exp^] and others, would be high priority to complete similar intron-retention studies, as chemical probes have already been developed [31, 36-38]. Clearly, such studies should be completed in patient-derived cell lines whenever possible.

Collectively, these studies show that there are a great many ways to affect RNA function that can be leveraged for therapeutic development. Further, systems that model native processing will be required to understand and capture the full potential of compounds to target RNA and rescue disease biology. The development of chemical probes and lead medicines targeting structured RNA, particularly human RNAs, are in their infancy. It will be interesting to see the types of chemical structures, their features, and the functional responses that emerge.

## Acknowledgements

We dedicate this work to our inspirational colleague Prof. K. Barry Sharpless on the occasion of being awarded the Priestly Medal. We thank Jessica Childs-Disney for help writing this manuscript and the agencies that funded this work including the National Institutes of Health (DP1-NS096898 to MDD and F31-NS110269-02 to AJA), the Muscular Dystrophy Association (grant 380467 to MDD), and a Fullbright Fellowship (to RIB).

