## Supporting Methods for "A Toxic RNA Catalyzes the Cellular Synthesis of Its Own Inhibitor, Shunting It to Endogenous Decay Pathways"

Table of Contents:

I. Supplementary Table and Figures (S1-S15)
II. Experimental Procedures
III. Compound synthesis
IV. References

### I. Supplementary Table & Figures

**Table S1.** Sequences of primers used for PCR

| Gene | Forward Primer (5'-3') | Reverse Primer (5'-3') | Purpose |
|---|---|---|---|
| *MAP4K4* | CCTCATCCAGTGAGGAGTCG | TGGTGGGAGAAATGCTGTATGC | RT-PCR |
| *IR* | CCAAAGACAGACTCTCAGAT | AACATCGCCAAGGGACCTGC | RT-PCR |
| *CNBP* i1 3'' SS | GAACTTTCAGTGGTTTAATGCTG | CCGCAGTTATAGCAGGCTTC | RT-PCR |
| *GAPDH* | AAGGTGAAGGTCGGAGTCAA | AATGAAGGGGTCATTGATGG | qPCR |
| *CNBP* Intron1 | ATTCCAAGGTTGGTTGAAGC | AACCCAAACCAATGAAGCTG | qPCR |
| *CNBP* mature mRNA | AAACTGGTCATGTAGCCATCAAC | AATTGTGCATTCCCGTGCAAG | qPCR |
| *MBNL1* | TTCATCCACCCCCACATTTA | TTGGCTAGTTGCATTTGCTG | qPCR |

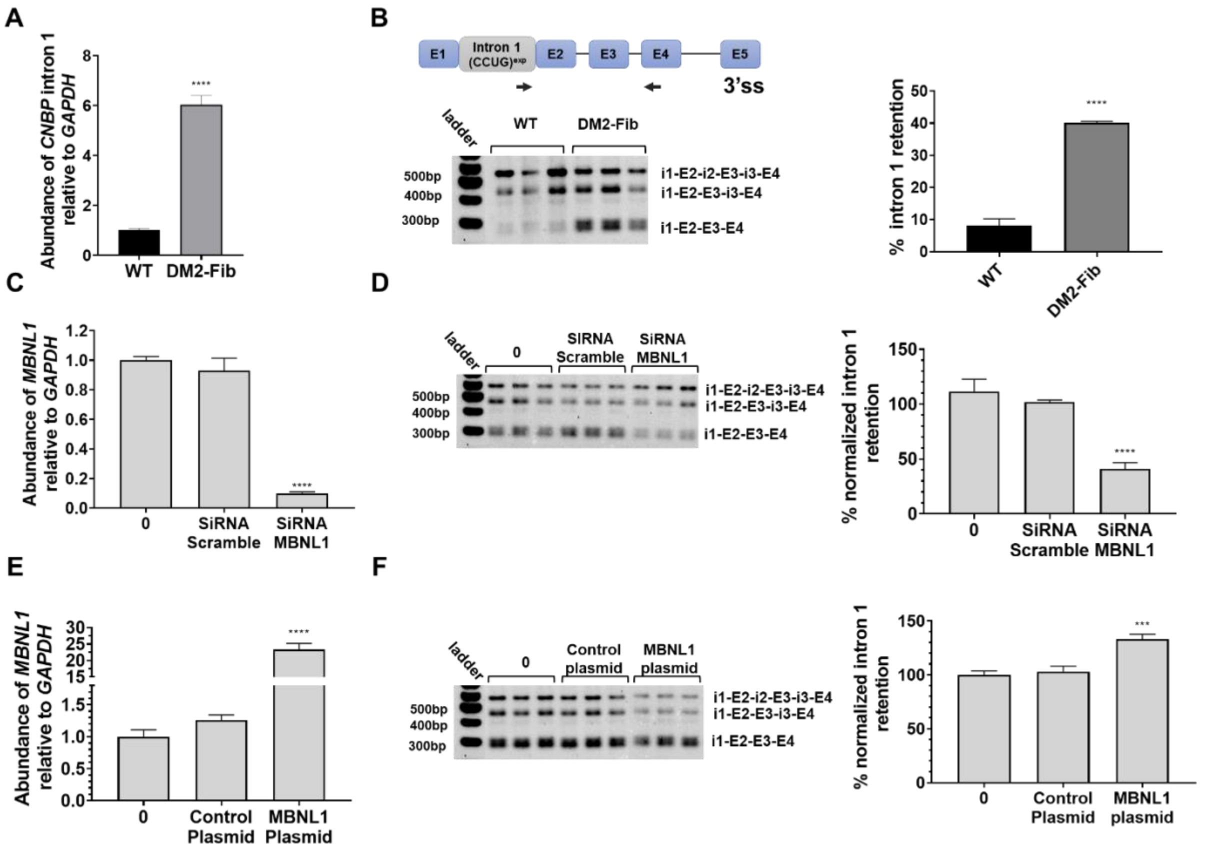


**Figure S1:** *MBNL1*'s effect on *CNBP*-retained species in DM2 patient-derived fibroblasts. (*A*) Abundance of *CNBP* intron 1 in WT fibroblasts compared to DM2 fibroblasts, as evaluated by RT-qPCR. (*B*) *CNBP* intron 1 retention levels in WT fibroblasts compared to DM2 fibroblasts, as evaluated by RT-PCR (left) and its quantification (right). Schematic of the *CNBP* i1 3′ss (2 primers - arrows) RT-PCR assay. *i* indicates intron and *E* indicates Exon. (*C, D*) Effect of MBNL1 knock-out on *CNBP* isoforms using siRNA in DM2 fibroblasts. (*C)* Relative abundance of *MBNL1* measured by RT-qPCR. (*D*) Representative gel image of *CNBP* intron 1 retention levels as measured by RT-PCR (left) and its quantification (right). (*E, F*) MBNL1 knock-in using a plasmid encoding MBNL1 in DM2 fibroblasts. (*E*) Relative abundance of *MBNL1* as measured by RT-qPCR. (*F*) Representative gel image of the effect of MBNL1 knock-in on *CNBP* isoforms using a transfected MBNL1 plasmid, as measured by RT-PCR (left) and its quantification (right). Error bars represent SD. ***P* < 0.01, ****P* < 0.001, *****P* < 0.0001 as determined by a one-way ANOVA (n = 3)

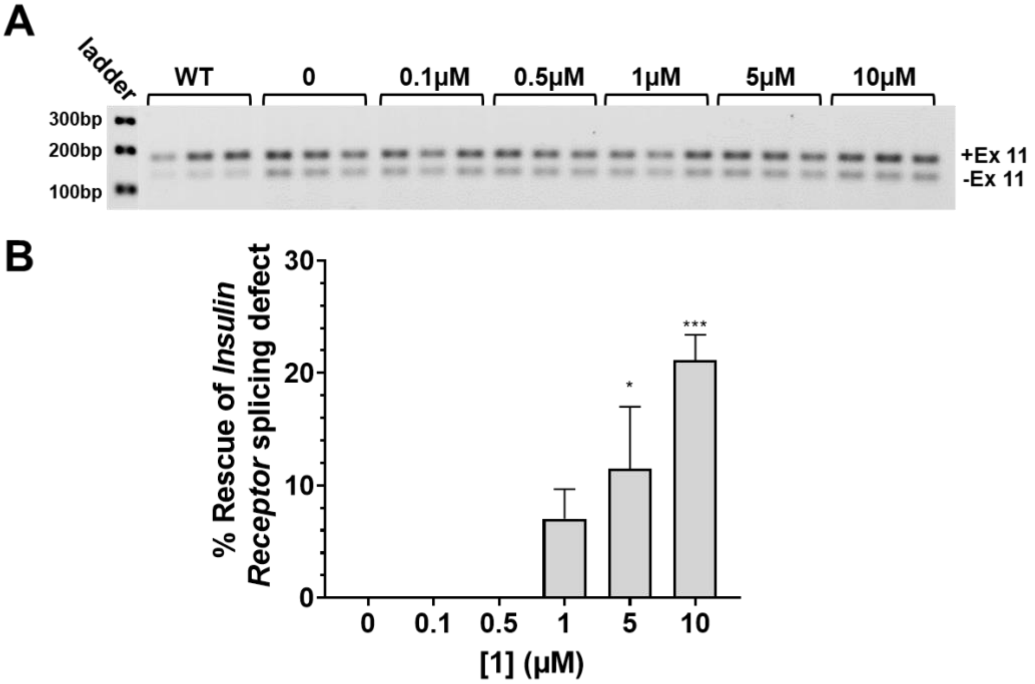


**Figure S2:** Ability of **1** to rescue aberrant splicing events in DM2 patient-derived fibroblasts. (*A*) Representative gel image of the alternative splicing of *Insulin Receptor* (*IR*) exon 11 in DM2 fibroblasts treated with **1**. (*B*) Quantification of RT-PCR analysis of the *IR* exon 11 splicing defect treated with **1**. Error bars represent SD. **P* < 0.5; ****P* < 0.001; as determined by a one-way ANOVA (n = 3).

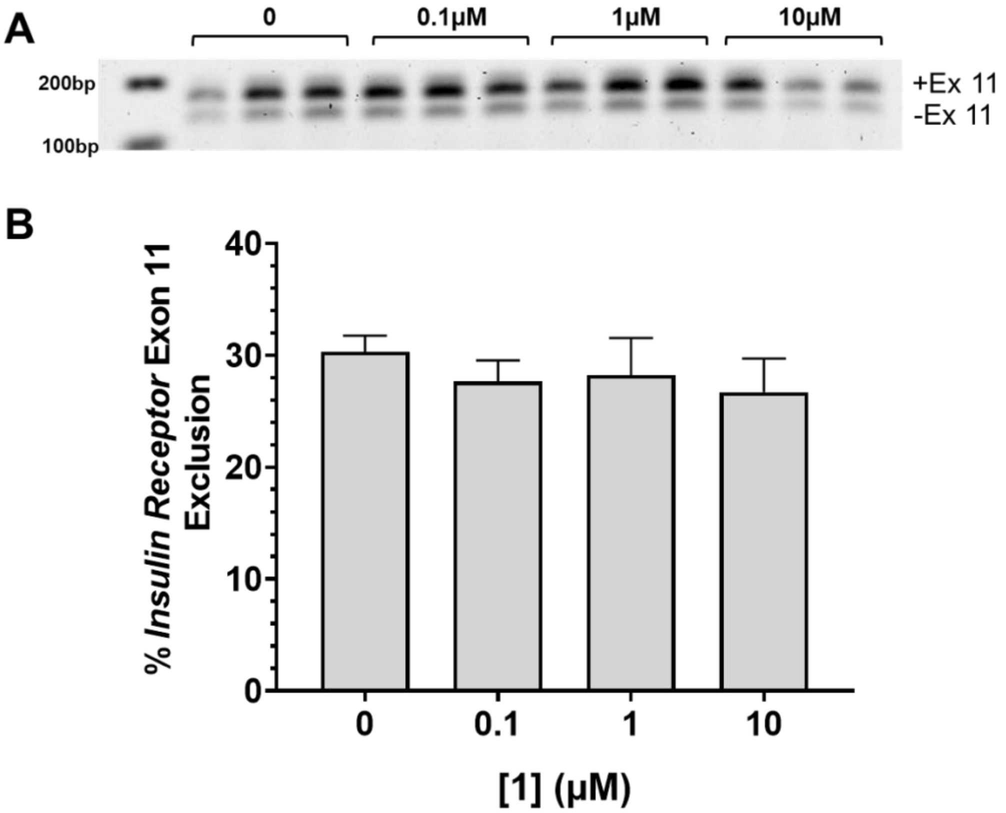


**Figure S3:** Evaluation of **1** in healthy fibroblasts. (*B*) Representative gel image of *IR* exon 11 splicing in WT fibroblasts treated with **2**. (*C*) Quantification of *IR* exon 11 splicing defect. Error bars represent SD (n=3).

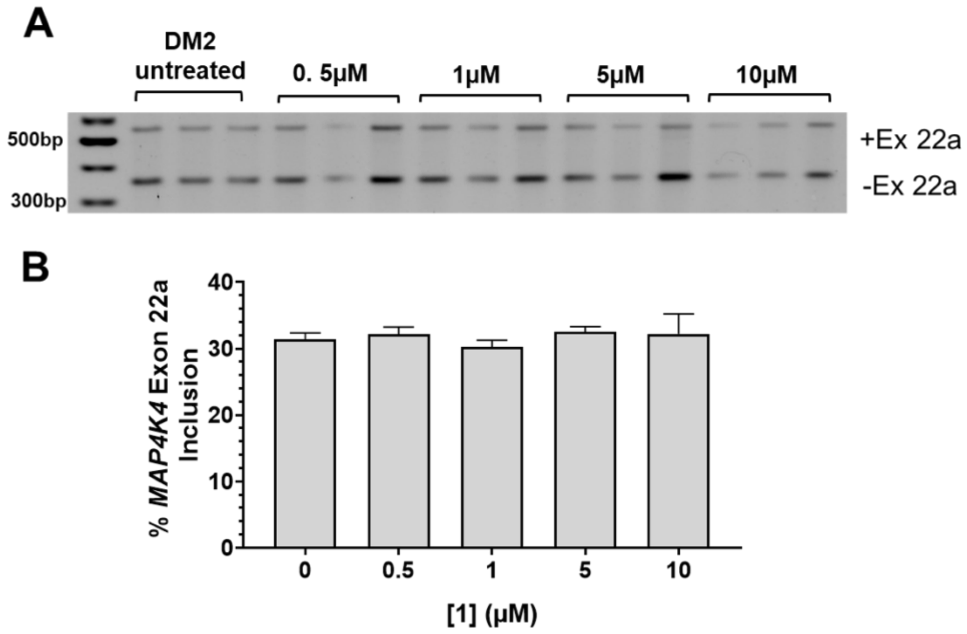


**Figure S4:** Evaluation of **1** in DM2 fibroblasts. (*A*) Representative gel image of *MAP4K4* exon 22a alternative splicing (non-MBNL1 regulated) in DM2 fibroblasts treated with **1**. (*B*) Quantification of *MAP4K4* exon 22a splicing. Error bars represent SD (n=3).

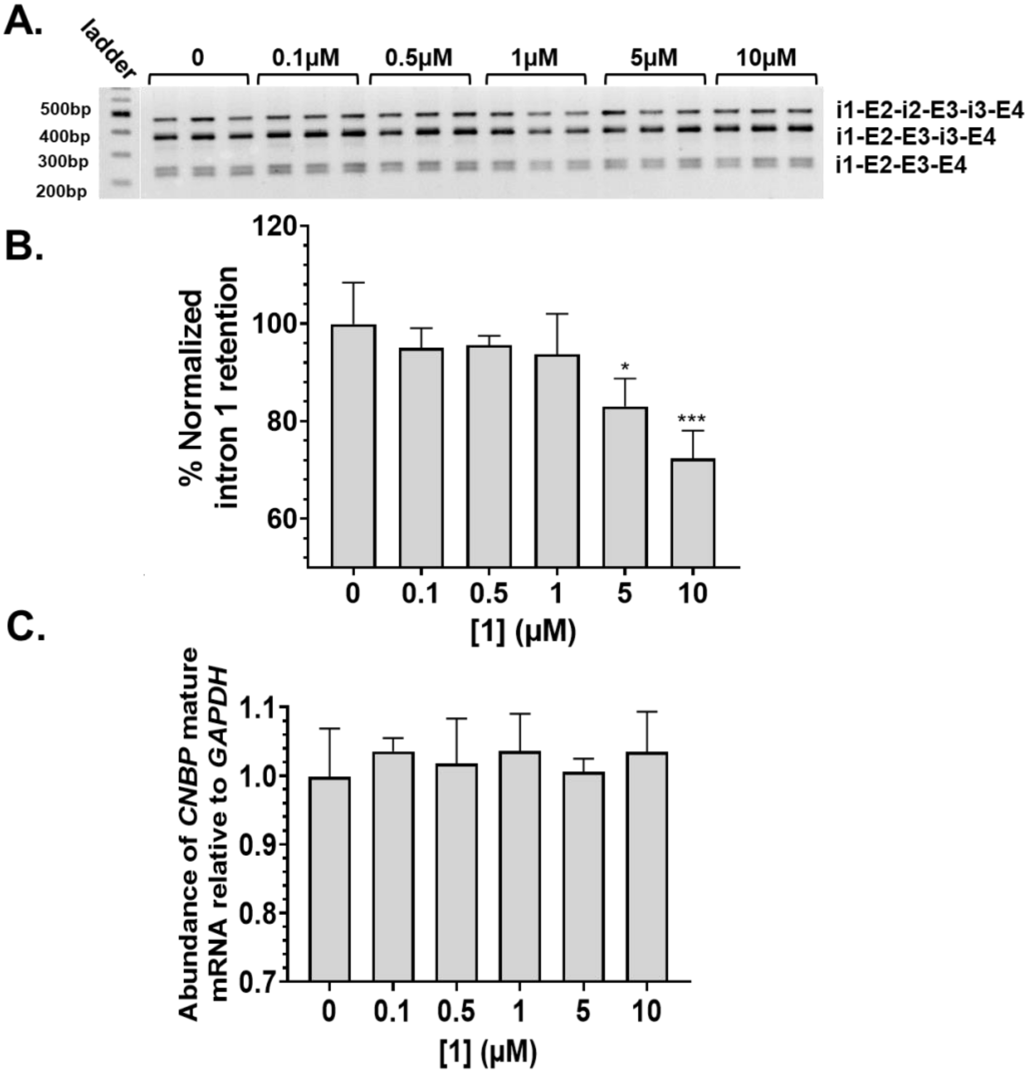


**Figure S5:** Ability of **1** to rescue aberrant *CNBP* splicing in DM2 patient-derived fibroblasts. (*A*) Representative gel image of *CNBP* intron 1 retention in DM2 fibroblast treated with **1**. (*B*) Quantification of RT-PCR analysis of *CNBP* intron 1 retention treated with **1**. (*C*) Effect of **1** on abundance of *CNBP* mature mRNA. Error bars represent SD. *$P < 0.5$; ***$P < 0.001$, as determined by a one-way ANOVA (n = 3)

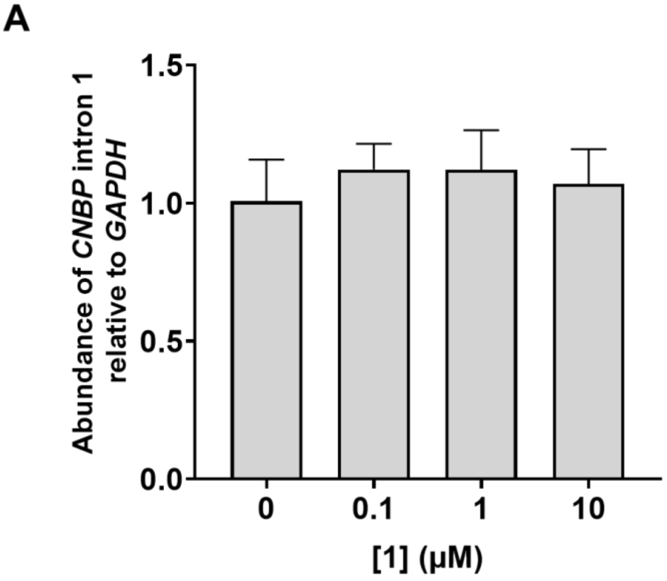


**Figure S6:** Evaluation of **1** in healthy fibroblasts. (*A*) RT-qPCR analysis of *CNBP* intron 1 abundance in WT fibroblasts treated with **2**. Error bars represent SD (n=3).

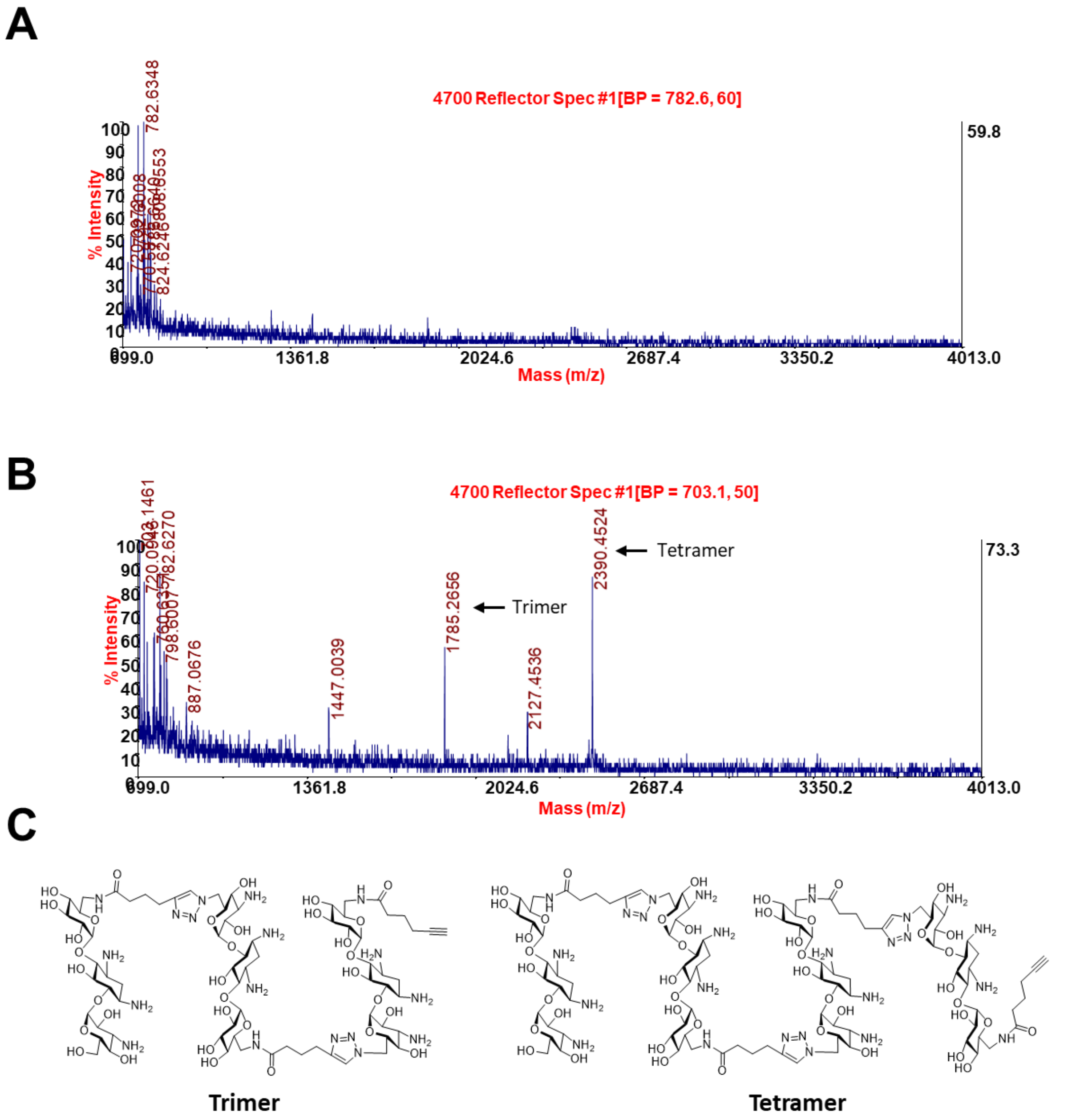


**Figure S7**. Results of mass spectral analysis of *in cellulis* click reactions between **2** and **3** in DM2 and healthy fibroblasts. (*A*) MALDI MS spectrum of compounds isolated from healthy fibroblast cells. (*B*) MALDI MS spectrum of compounds isolated from DM2 fibroblasts, showing that a trimer and tetramer were formed. (*C*) Chemical structures of the *in cellulis* click products (trimer and tetramer).

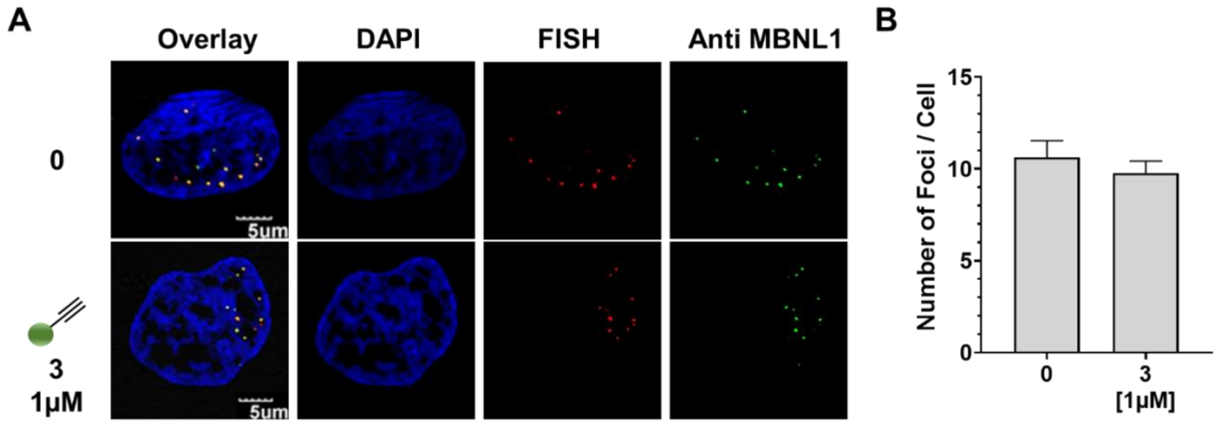


**Figure S8:** Evaluation of **3** in DM2 fibroblasts. (*A*) Representative images of r(CCUG)$^{exp}$-MBNL1 foci in DM2 fibroblasts treated with **3**. (*B*) Quantification of r(CCUG)$^{exp}$-MBNL1 foci/cell. Error bars represent SD (n=3; biological replicates, 40 nuclei counted per replicate).

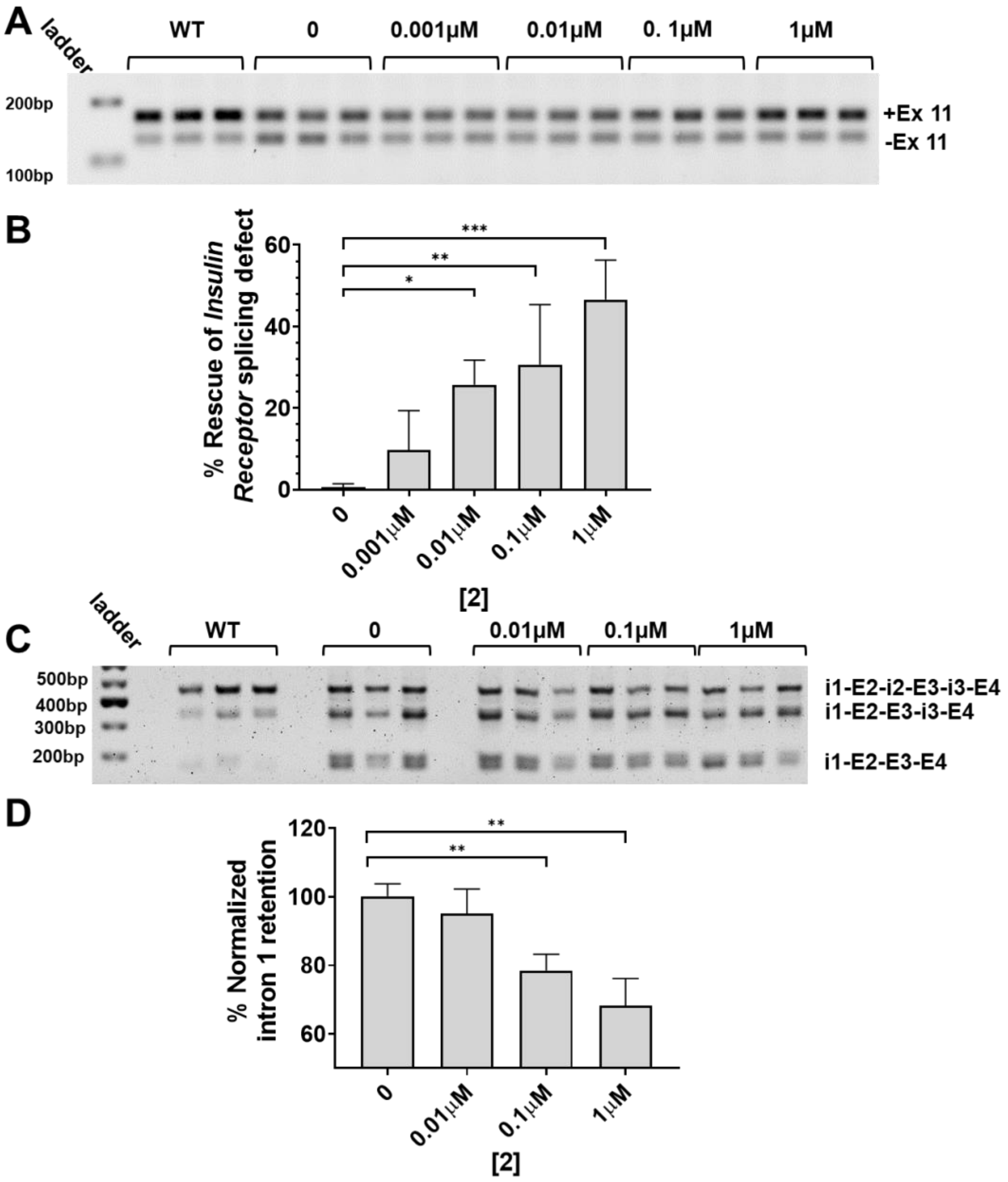


**Figure S9:** Ability of **2** to rescue aberrant splicing events in DM2 fibroblasts. (*A*) Representative gel image of *IR* exon 11 alternative splicing in DM2 fibroblasts treated with **2**. (*B*) Quantification of RT-PCR analysis of the *IR* exon 11 splicing defect treated with **2**. (*C*) Representative gel image of *CNBP* intron 1 retention in DM2 fibroblasts treated with **2**. (*D*) Quantification of RT-PCR analysis of *CNBP* intron 1 retention in cells treated with **2**. Error bars represent SD. **P* < 0.5, ***P* < 0.01, ****P* < 0.001 as determined by a one-way ANOVA (n = 3).

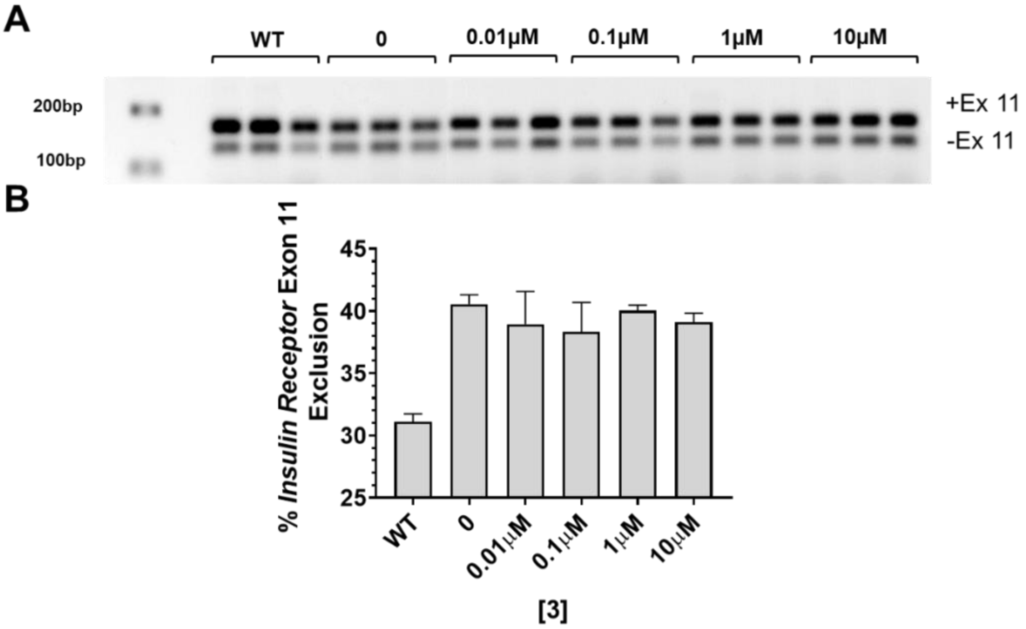


**Figure S10:** Evaluation of **3** in DM2 fibroblasts. (*A*) Representative gel image of *IR* exon 11 alternative splicing in DM2 fibroblast treated with **3**. (*B*) Quantification of *IR* exon 11 splicing defect. Error bars represent SD (n=3; biological replicates).

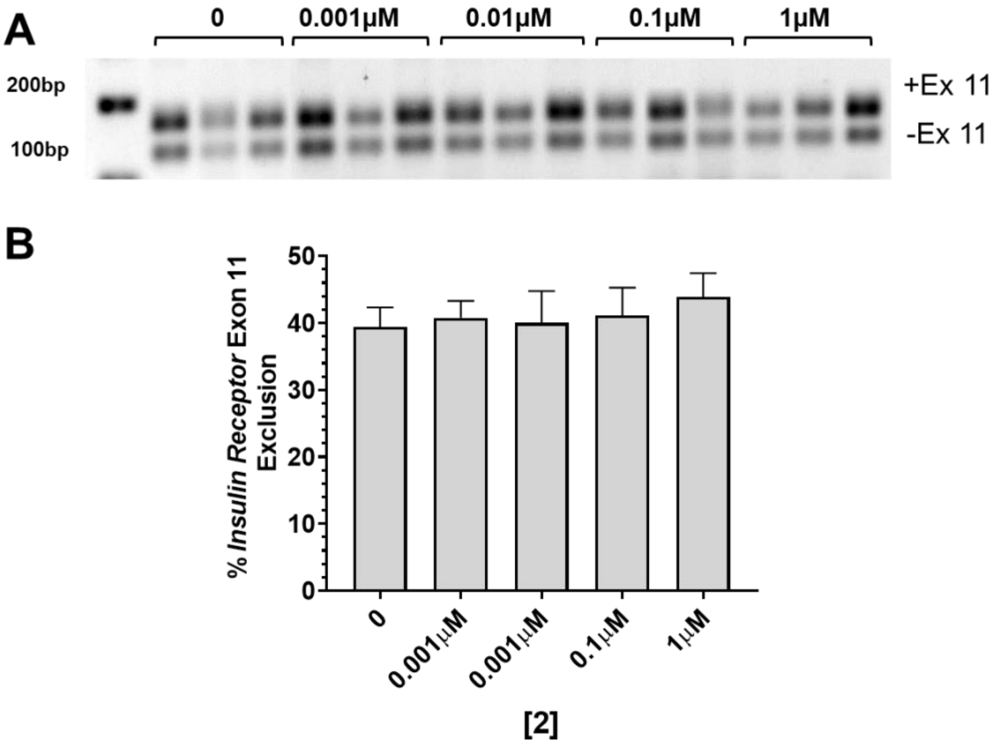


**Figure S11:** Evaluation of **2** in healthy (WT) fibroblasts. (*A*) Representative gel image of *IR* exon 11 alternative splicing in WT fibroblasts treated with **2**. (*B*) Quantification of *IR* exon 11 splicing defect. Error bars represent SD (n=3).

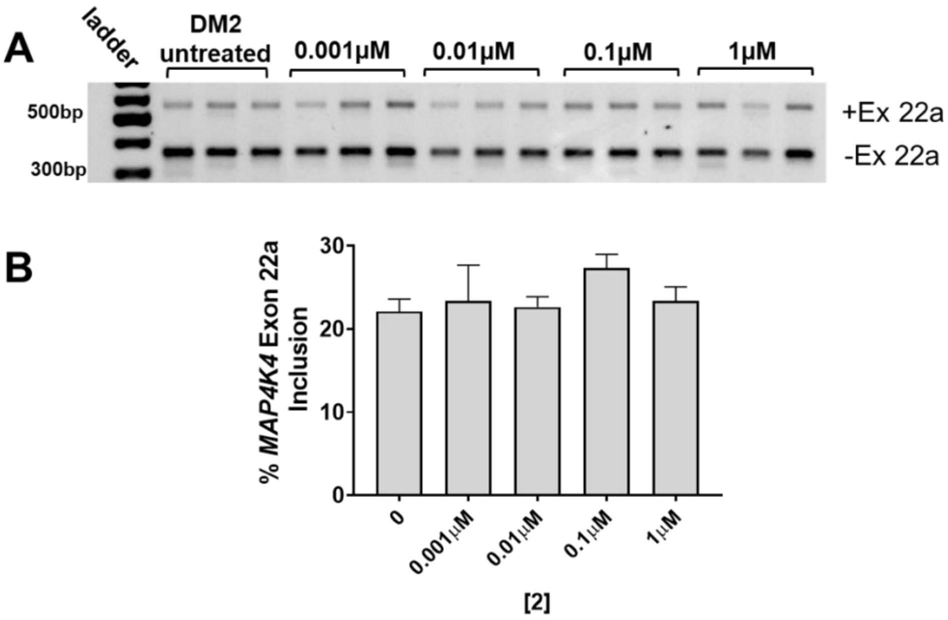


**Figure S12:** Evaluation of **2** in DM2 fibroblasts. (*A*) Representative gel image of *MAP4K4* exon 22a alternative splicing (non-MBNL1 regulated) treated with **2**. (*B*) Quantification of *MAP4K4* exon 22a splicing. Error bars represent SD (n=3).

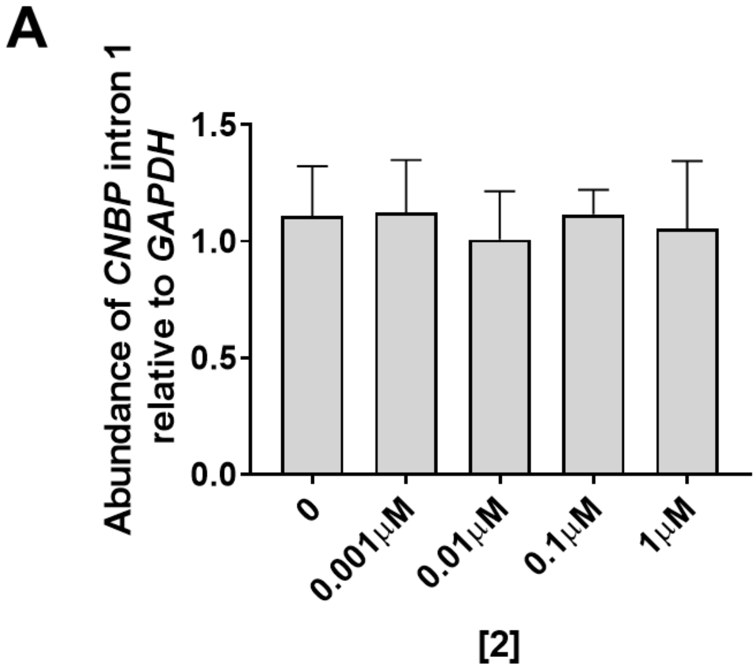


**Figure S13:** Evaluation of **2** in healthy (WT) fibroblasts. (*A*) RT-qPCR analysis of *CNBP* intron 1 abundance in WT fibroblasts treated with **2**. Error bars represent SD (n=3).

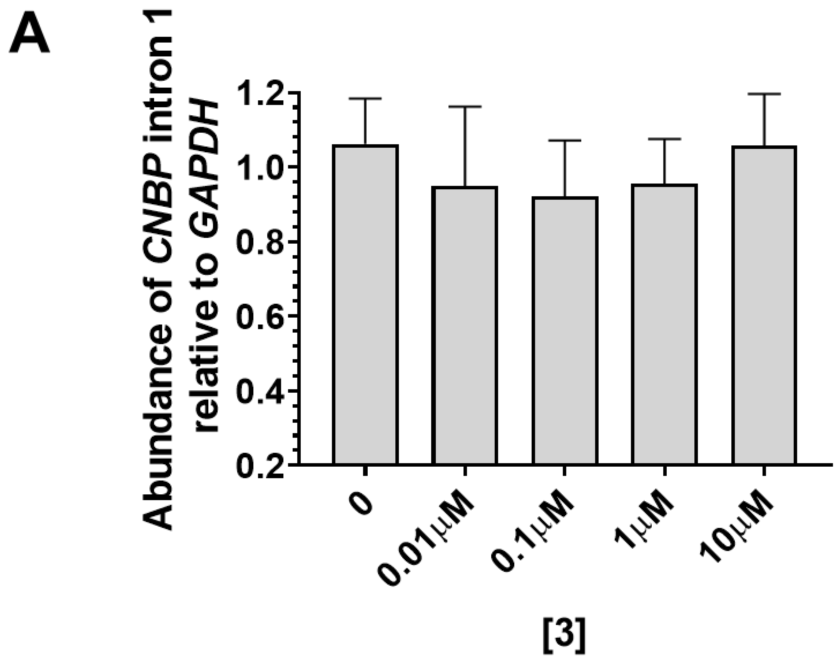


**Figure S14:** Evaluation of **3** in DM2 fibroblasts. (*A*) RT-qPCR analysis of *CNBP* intron 1 abundance in DM2 fibroblasts treated with **3**. Error bars represent SD (n=3).

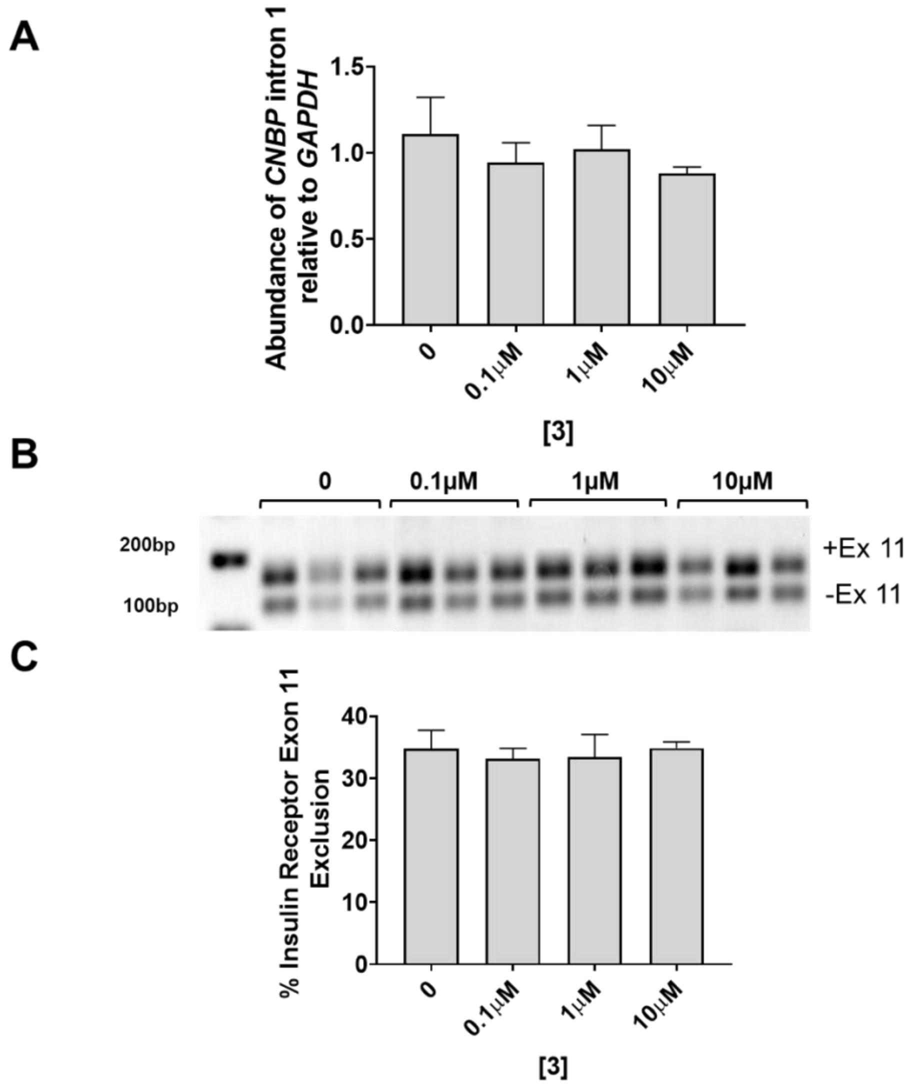


**Figure S15:** Evaluation of **3** in healthy fibroblasts. (*A*) RT-qPCR analysis of *CNBP* intron 1 abundance in WT fibroblasts treated with **3**. (*B*) Representative gel image of *IR* exon 11 alternative splicing in WT fibroblasts treated with **3**. (*C*) Quantification of *IR* exon 11 splicing defect. Error bars represent SD (n=3).

### II. Experimental Procedures

**Cell lines:** Compounds were tested in two cell lines: (i) a DM2 patient-derived fibroblasts (DM11; Center for NeuroGenetics); and (ii) fibroblasts from a healthy donor (wild type; WT GM07492; Coriell Institute).

**Cell culture:** All cells were maintained at 37 °C with 5% $CO_2$. DM2 fibroblasts were cultured in DMEM / High glucose (HyClone) supplemented with 20% FBS and 1% Antibiotic-Antimycotic (Corning). WT fibroblasts were cultured in MEM (Corning) supplemented with 10% FBS (Sigma), 1% Glutagro (Corning), and 1% Antibiotic-Antimycotic (Corning).

Cells were transfected with EGFP (empty vector control) and MBNL1 plasmids (to overexpress MBNL1) in 6-well plates with Lipofectamine 3000 per the manufacturer's protocol. After 5 h, the medium was replaced with growth medium, and the cells were grown for an additional 24 h. siRNA-MBNL1 (Sense:5'-C.A.C.U.G.G.A.A.G.U.A.U.G.U.A.G.A.G.A.dT.dT-3'; Antisense: 5'-U.C.U.C.U.A.C.A.U.A.C.U.U.C.C.A.G.U.G.dT.dT-3' purchased from Dhamacon), and siRNA control (purchased from Santa S-39 Cruz Biotechnology; control siRNA catalog #: sc-37007) were transfected into cells using Lipofectamine RNAiMAX Reagent per the manufacturer's protocol for 48 h.

Cells were treated with compound **1**, **2** and **3** at concentrations ranging from 10μM-0.1μM, 1μM-0.1μM and 10μM-0.01μM respectively, in growth medium for 48 h at 37 °C with 5% $CO_2$.

**Analysis of abundance of CCUG-containing transcripts:** Cells were grown in 6-well plates and treated or transfected as described in "**Cell Culture**". After 48 h, the cells were lysed, and total RNA was harvested using a Zymo Quick RNA miniprep kit. Approximately 1 μg of total RNA was reverse transcribed using a qScript cDNA synthesis kit (20 μL total reaction volume, Quanta BioSciences). A 2 μL aliquot of the RT reaction was used for each primer pair for qPCR, which was completed with SYBR Green Master Mix on an Applied Biosystems 7900HT Fast Real-Time PCR System. Relative abundance of each transcript was calculated by normalizing to *GAPDH*.

**Evaluation of nuclear foci using fluorescence *in situ* hybridization (FISH):** FISH was used to determine the small molecules' effects on formation and disruption of nuclear foci as previously described.(1) Briefly, DM2 fibroblasts were grown to ~40% confluence in a Mat-Tek 96-well glass bottom plate in growth medium. Cells were then treated with **1**, **2** or **3** for 48 h. To fix the cells, the compound-containing growth medium was removed, the cells were washed with 1× DPBS, and 100 μL of 4% formaldehyde in 1× DPBS was added. After incubating at 37°C for 10 min, the cells were washed five times with 1× DPBS at 37°C for 2 min and then twice with 100 μL of 0.1% Triton X-100 in 1× DPBS for 5 min at 37°C. Next, 100 μL of 30% formamide in 2× SSC buffer was added, and the cells were incubated for 10 min at room temperature. The FISH probe, 5'-Cy3-$(CAGG)_{10}$ (140 nM; purchased from Integrated DNA Technologies, Inc.) was added to each well in 2× SSC containing 30% formamide, 2 μg/μL BSA, 1 μg/μL yeast tRNA, and 2 mM vanadyl complex), and the cells were incubated at 37 °C overnight. The cells were then washed with 100 μL of 2× SSC containing 30% formamide at 37 °C for 30 min and then 100 μL of 2× SSC buffer for 30 min at 37°C. For MBNL1 immunostaining, 20 μL of 1:5 anti-MBNL1 in 2× SSC was added, and the cells were incubated at 37 °C for 1 h. The cells were washed three times with 100 μL of 1× DPBS containing 0.1% Triton X-100 for 5 min at 37 °C and then incubated with 1:200 dilution of anti-mouse IgG Dylight 488 in 2× SSC at 37°C for 1 h. The cells were then washed three times with 100 μL of 1× DPBS containing 0.1% Triton X-100 for 5 min at 37 °C and then with 1× DPBS for 5 min at 37°C. Finally, 100 µL of 1 µg/mL DAPI was added, and cells were incubated for 5 min at 37 °C. After washing with 1× DPBS twice, the cells were placed in 100 µL of 1× DPBS and imaged using an Olympus Fluoview 1000 confocal microscope at 100× magnification.

**Evaluation of pre-mRNA splicing:** Cells were grown in 6-well plates and treated as described in "**Cell Culture**". After 48 h, the cells were lysed, and total RNA was harvested using a Zymo Quick RNA miniprep kit. Approximately 1 µg of total RNA was reverse transcribed using a qScript cDNA synthesis kit (20 µL total reaction volume, Quanta BioSciences); 2 µL of the RT reaction was used for PCR amplification using GoTaq DNA polymerase (Promega). RT-PCR products were observed after 30 cycles of: 95 °C for 30 s; 58°C for 30 s; and 72°C for 1 min followed by an additional extension at 72 °C for 5 min. Products were separated on a 2% agarose gel (110 V for 1 h in 1× TBE buffer), visualized by staining with ethidium bromide, and imaged using a Typhoon 9410 variable mode imager. Gels were quantified using ImageJ. Percent rescue was calculated by dividing the difference between treated and untreated DM2 samples by the difference between untreated DM2 and WT samples (Equation 1).

$$\% \, Rescue = \frac{\% \, exon \; inclusion \; DM2 - \% \, exon \; inclusion \; treated}{\% \, exon \; inclusion \; DM2 - \% \, exon \; inclusion \; WT} * 100 \qquad \text{(Eq. 1)}$$

**Identification of *in cellulis* clicked products by mass spectrometry.** DM2 fibroblasts were grown in 100 mm dishes in growth medium. Cells were treated with 25 µM of **2** and/or 25 µM of **3**. Compound **3** was added to limit the molecular weight of the oligomeric products in order to enable detection by LC-MS. After 48 h, the cells were scraped from the plate and lysed by freezing and thawing in 10% water in acetonitrile. The thawed lysate was concentrated and re-suspended in 1 mL of 10% water in acetonitrile. Insoluble cellular debris was pelleted, and the supernatant was used for mass spectral analysis. Mass spectrometry was performed with an Applied Biosystems MALDI ToF/ToF Analyzer 4800 Plus using an α-cyano-4-hydroxycinnamic acid matrix.

### III. Compounds synthesis:

Compounds **1, 2** and **3** were synthesized as previously described and dissolved in nanopure water. (2, 3)
